# Peripheral Nerve Magnetoneurography with Optically Pumped Magnetometers

**DOI:** 10.1101/2021.05.18.444539

**Authors:** Yifeng Bu, Jacob Prince, Hamed Mojtahed, Donald Kimball, Vishal Shah, Todd Coleman, Mahasweta Sarkar, Ramesh Rao, Mingxiong Huang, Peter Schwindt, Amir Borna, Imanuel Lerman

## Abstract

Electrodiagnosis is routinely integrated into clinical neurophysiology practice for peripheral nerve disease diagnoses such as neuropathy, demyelinating disorders, nerve entrapment/impingement, plexopathy or radiculopathy. Measured with conventional surface electrodes, the propagation of peripheral nerve action potentials along a nerve is the result of ionic current flow which, according to Ampere’s Law, generates a small magnetic field that is also detected as an “action current” by magnetometers such as superconducting quantum interference device (SQUID) Magnetoencephalography (MEG) systems. Optically pumped magnetometers (OPMs) are an emerging class of quantum magnetic sensors with a demonstrated sensitivity at the 1 fT/√Hz level, capable of cortical action current detection. But OPMs were ostensibly constrained to low bandwidth therefore precluding their use in peripheral nerve electrodiagnosis. With careful OPM bandwidth characterization, we hypothesized OPMs may also detect compound action current signatures consistent with both Sensory Nerve Action Potential (SNAP) and the Hoffmann Reflex (H-Reflex). In as much, our work confirms OPMs enabled with expanded bandwidth can detect the magnetic signature of both the SNAP and H-Reflex. Taken together, OPMs now show potential as an emerging electrodiagnostic tool.

## 1 Introduction

The human peripheral nervous system is composed of an intricate network of motor, sensory and autonomic neural structures (Barr, 1974;Sulaiman et al., 2001;Mai and Paxinos, 2011) that if injured can result in peripheral neuropathic pain. Electrodiagnosis augments the clinicians’ ability to gauge the severity of the neuropathy, and the distribution of neuropathic dysfunction, that consequently guide clinical pharmacologic, minimally invasive or invasive surgical interventions (Do Campo, 2021). Well established in clinical neurophysiology practice, conventional (surface electrode) electrodiagnostic, routinely identify peripheral neuropathy, and peripheral nerve entrapment with the Sensory Nerve Action Potential (SNAP); whereas plexopathy, demyelinating disease and radiculopathy are identified with the Hoffmann Reflex (H-Reflex). SNAP measures electrically evoked sensory nerve fiber action potentials of the upper and lower extremities; while the H-Reflex, an electrically evoked sensory-motor mono-synaptic spinal reflex, may be recorded from any surface muscle with an accessible nerve (Burke et al., 1989; Pierrot-Deseilligny and Burke, 2005). Abnormal (delayed 3-4 ms) H-Reflexes are known to occur at the onset of nerve root compression at the very early stage of nerve root injury even when imaging is normal and remain abnormal until the compression ceases (i.e., with selective nerve root injection or corrective surgical intervention)(Jin et al., 2010;Zheng et al., 2014). Further H-Reflexes are absent in acute inflammatory demyelinating polyneuropathy (Guillain–Barré syndrome)(Bamford and Davis, 2020). Abnormal SNAP measures (delay or amplitude decrement) are regularly identified early in peripheral nerve entrapment and are pathognomonic in peripheral neuropathy (Krarup, 2004;Yang et al., 2021). Although surface electrode SNAP and H-Reflexes are proven integral electrodiagnostic measures; novel nascent magnetoneurography techniques are in development, capable of compound action current measurement in the peripheral nerve and muscle.

Super conducting quantum interference devices (SQUID) sensor arrays are a non-invasive diagnostic employed to detect cortical neuronal action potentials with magnetoencephalography (MEG). Peripheral nerve action currents in the human upper extremity (median and ulnar nerve) (termed magnetoneurography) (Hari et al., 1989;Hashimoto et al., 1994;Lang et al., 1998) and, more recently, in the dorsal root ganglion and spinal cord (termed magnetospinography) have successfully been detected with SQUID sensor arrays (Adachi et al., 2009). However, conventional SQUID sensor use is highly limited, as they require cryogenic temperature provided with liquid helium (4 K or -269 °C) and magnetically shielded rooms (MSRs). Moreover, SQUID sensors are housed in a rigid Dewar which prohibit adjacent placement to the sensory, motor nerve or the target muscle that inherently requires highly mobile and conformal sensor arrays.

QuSpin OPM sensors are conformal with a 6.5 mm sensor stand-off that in aggregate contribute to an improved magnetic signal-to-noise ratio therefore capable of surpassing current SQUID sensor technology due to closer proximity to the neuronal source (Iivanainen et al., 2017;Hill et al., 2020). Recently Broser et.al., (Broser et al., 2018;Broser et al., 2020) showed that commercial Gen-1 OPMs (QuSpin Inc, CO, USA) detect compound muscle action currents consistent with motor neuron activation. Expounding on this work, we employed the commercially available Gen-2 OPM (QuSpin Inc, CO, USA) with the aim to measure upper extremity action currents equivalent to the SNAP and H-Reflex. To achieve this aim we meticulously: 1) characterized the frequency response, phase response, and sensitivity of the OPM sensor, and 2) validated the temporal resolution of the OPM’s recording of upper extremity action currents (equivalents of SNAP and H-Reflex) when equated to conventional surface electrodes. In concert, these results demonstrate preliminary potential for OPM as a peripheral nerve electrodiagnostic tool.

## 2 Methods

### 2.1 OPM Operational Bandwidth

QuSpin Gen2 OPMs were used to measure median nerve compound action currents. In comparison to the Gen-1 OPM (dimensions: 13.0×19.0×110.0 mm^3^), Gen-2 OPMs have a significantly smaller footprint (12.4×16.6×24.4 mm^3^) that improves conformal placement of the OPM array. Gen-2 OPMs demonstrate improved sensitivity (7-10 fT/√Hz in ideal conditions) and have an extended tolerance to background magnetic field (up to 200 nT) while maintaining ±5 nT dynamic range (Shah and Romalis, 2009). The OPM integrates a sixth-order digital filter at 500 Hz to eliminate residual response above this frequency.

QuSpin OPM sensor response capability of magnetic field frequencies (up to 500 Hz) was identified with response to a frequency chirp function. The voltage chirp function (swept linearly from 1 Hz to 700 Hz over a 2 second period) was converted into a proportional magnetic chirp by a copper-wire Helmholtz Coil (two 7.5 cm radius loops with 5 coils per loop at 7.5 cm separation) and associated current supply circuit (**Supplemental Figure 1**). Based on electromagnetic theory, the amplitude of the current running through the Helmholtz coil is directly proportional to the amplitude of the magnetic field generated at the center of the two coils (Bronaugh, 1995;Wang et al., 2002;Cvetkovic and Cosic, 2007;de Melo et al., 2009). Accordingly, the current supply was controlled to generate a 600 pT peak-to-peak (pk-pk) alternating magnetic field at the OPM’s central location. Using the OPM recordings of repeated (N=11) chirp functions, the OPM magnitude and phase response was then calculated. Additionally, the sensitivity of the sensor was calculated by computing the power spectral density of an empty room recording and dividing this value by the normalized frequency response curve. Before the creation of the sensitivity plot, the OPM signal was filtered for powerline sources (i.e., 60 Hz and its harmonics) that allowed for removal of these the sensitivity spikes. Finally, using the same Helmholtz Coil setup with a varying input current amplitude, the OPM response to a 73 Hz sinusoidal magnetic field input was recorded at amplitudes measured between 15 and 1,500 pT. Magnetic recordings from this experiment were used to calculate the total harmonic distortion of the OPM at multiple measured signal amplitudes.

### 2.2 Experimental Setup and Procedures

The Institutional Review Board at the University of California San Diego (UCSD) Health Systems approved the experimental protocol (UCSD IRB:171154). Three healthy male subjects (age: 40±12 years) without history, signs, or symptoms of peripheral neuropathy, nerve entrapment syndrome or radiculopathy, gave their written consent. On the visit day, surface electrode and OPM-based peripheral nerve measurements were carried out within the UCSD Radiology Imaging Laboratory’s six-layer MSR (IMEDICO, Switzerland), to minimize the effects of powerline noise and the Earth’s magnetic field. The employed MSR has a shielding factor of 65-160 dB for the 0.01 Hz-10 Hz frequency range. Subjects were asked to remove all electronic equipment and metal accessories before entering the MSR to avoid magnetic noise and sensor saturation.

#### 2.2.1 Median Nerve Sensory Nerve Action Potential and Current Measure

The subject (N=1) was centered in the middle of the MSR in the supine position. A pillow was positioned below the subject’s left elbow to ensure comfortable yet maximal extension of the upper arm above the head. The median nerve SNAP and sensory nerve action current was measured between the brachialis and the biceps brachii. A 20 MHz portable ultrasound transducer (Butterfly IQ, Palo Alto, USA) was used to measure the median nerve depth, circumference, and area at both proximal bicep and distal bicep sites. Stimulating electrodes with cathode-anode distance of 1.5 cm were secured (with a Velcro strap) longitudinally over the left median nerve at the wrist with cathode proximal to anode. Two pairs of active and reference electrodes were localized (via ultrasound image guidance) over the median nerve separated by a 2 cm, 4 cm and 6 cm distance while the ground electrode was placed between the stimulator and the active electrode (**Figure 2A)**. In all measurements, the skin was carefully abraded with the Nuprep skin prep gel (Weaver and Company, CO, USA) and the EC3 Grass conductive adhesive gel (Natus, Pleasanton, CA, USA) was applied to each electrode cup with assurance of impedance maintenance at 5000 Ω or less. Subjects then underwent left median nerve supramaximal stimulation with the DS7AH constant stimulator (Digitimer Ltd, UK) using a 0.23 Hz, 500 μs pulse-width, monopolar square-wave.

**FIGURE 1.**
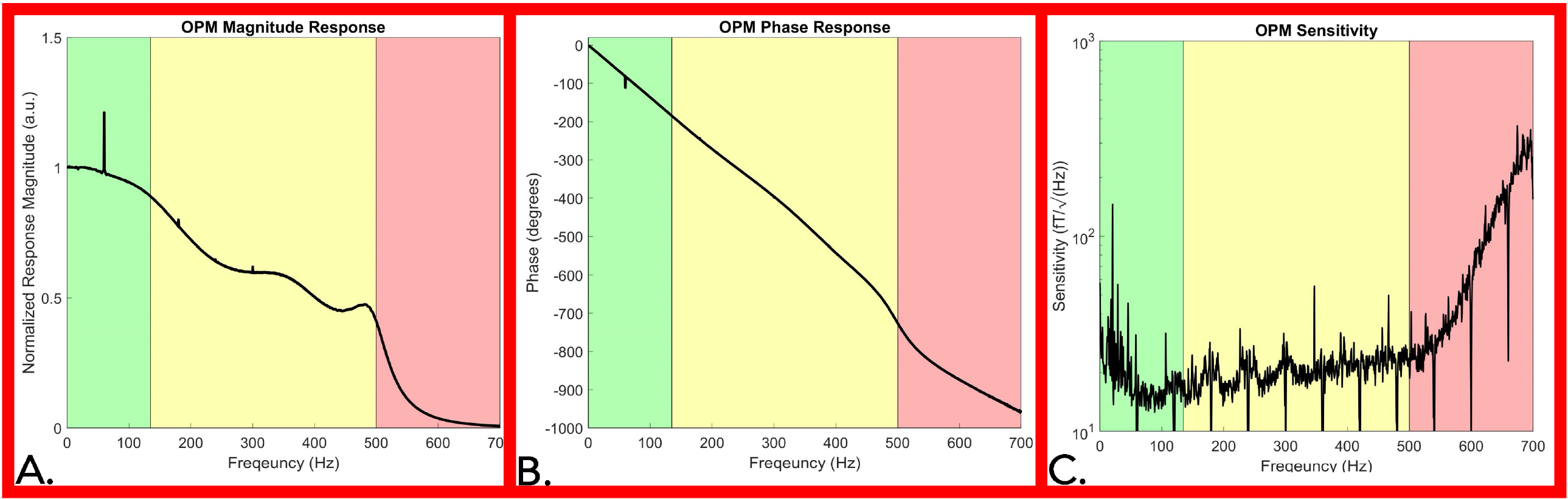
OPM frequency response and sensitivity characterization. **Panel A:** OPM magnitude response decreased as frequency increased, particularly above 135 Hz (**green shaded area**), while a reduced frequency response was recorded up to 500 Hz (**yellow shaded area**). Above 500 Hz a large reduction in response is noted as a direct effect of the Gen2 OPM hardware digital filter in the electronic module (**red shaded area**). **Panel B:** OPM Phase response is approximately linear (**green and yellow shaded area**) up to 500 Hz (**red shaded area). Panel C:** Sensitivity response of the OPM. OPM sensitivity was divided into three segments: below 135 Hz (**green shaded area)**, from 135 to 500 Hz (**yellow shaded area**), and from 500 to 700 Hz (**red shaded area**). A decrease in sensitivity is noted above 500 Hz. (OPM=Optically Pumped Magnetometer, Hz= Hertz, fT= femtotesla)

**FIGURE 2.**
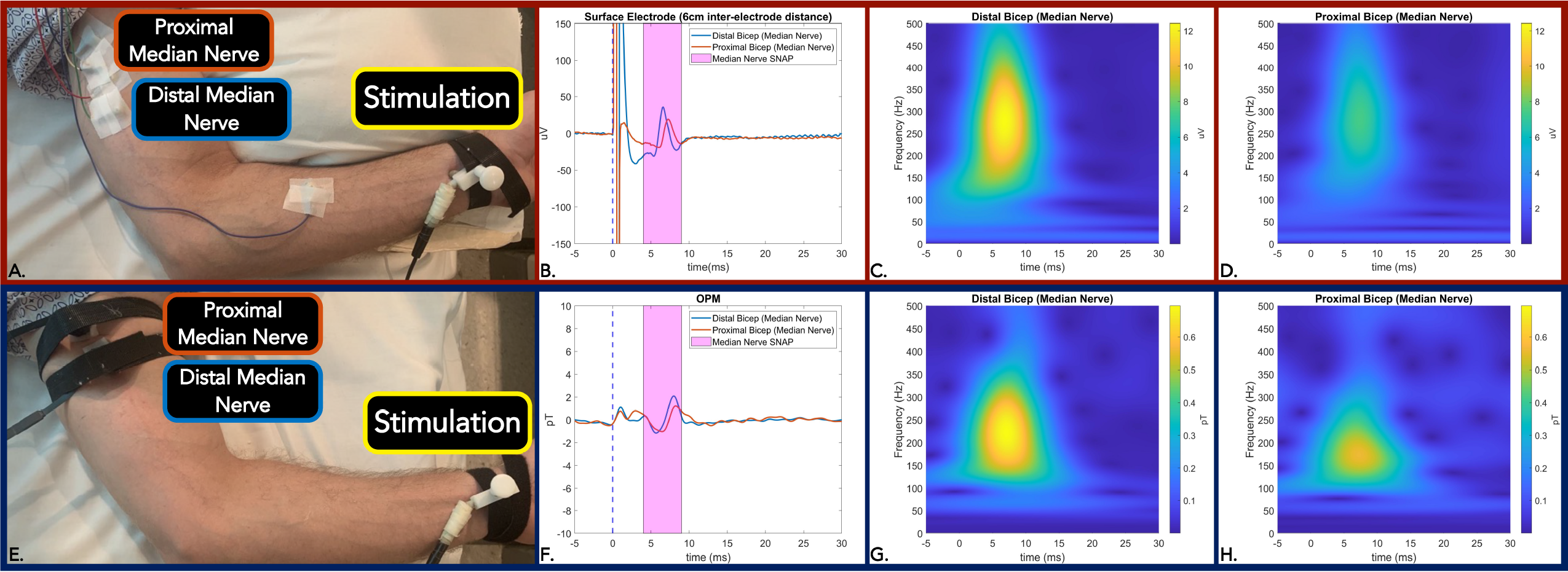
**Panel A, E:** Measurement setup of Median nerve SNAP with surface electrode **(A)** and equivalent action current by OPM **(E). Panel B and F:** Time-locked average comparison (from two sites; distal bicep=blue line and proximal bicep=red line) between surface electrode (6m interelectrode distance) **(B)** and OPM **(F)** demonstrate identical 0.8 ms temporal dispersion for both modalities. SNAP action potential/currents are marked in the magenta shaded area. **Panel C and D**: Surface electrode time-frequency analysis for SNAP measured by at distal bicep **(C)** and proximal bicep **(D). Panel G and H**: OPM time-frequency analysis for SNAP measured at distal bicep **(G)** and proximal bicep **(H)**. (OPM = Optically Pumped Magnetometers, SNAP= Sensory Nerve Action Potential, μV = microvolt, pT=picoTesla, ms=milliseconds)

#### 2.2.2 Median Nerve Hoffman Reflex

The H-reflex response was elicited in the Flexor Carpi Radialis muscle (FCR) by stimulator placement along the antecubital fossa longitudinal axis directly over the left median nerve on two subjects. The recording techniques employed for H-Reflex generally followed the methodologies of two studies (Christie et al., 2005;Khosrawi et al., 2015). Briefly, the active electrode was placed over the muscle belly of the FCR located one-third of the distance between the medial epicondyle and radial styloid while the reference electrode was placed over the FCR tendon insertion site at the wrist (**Figure 3A)**. Isometric contraction of the FCR muscle was used to facilitate the reflex response. Stimulation intensity was incrementally increased by 0.2 mA until a clear H-Reflex was elicited followed by amplitude decreased with subsequent maximal M-wave (Stowe et al., 2008).

**FIGURE 3.**
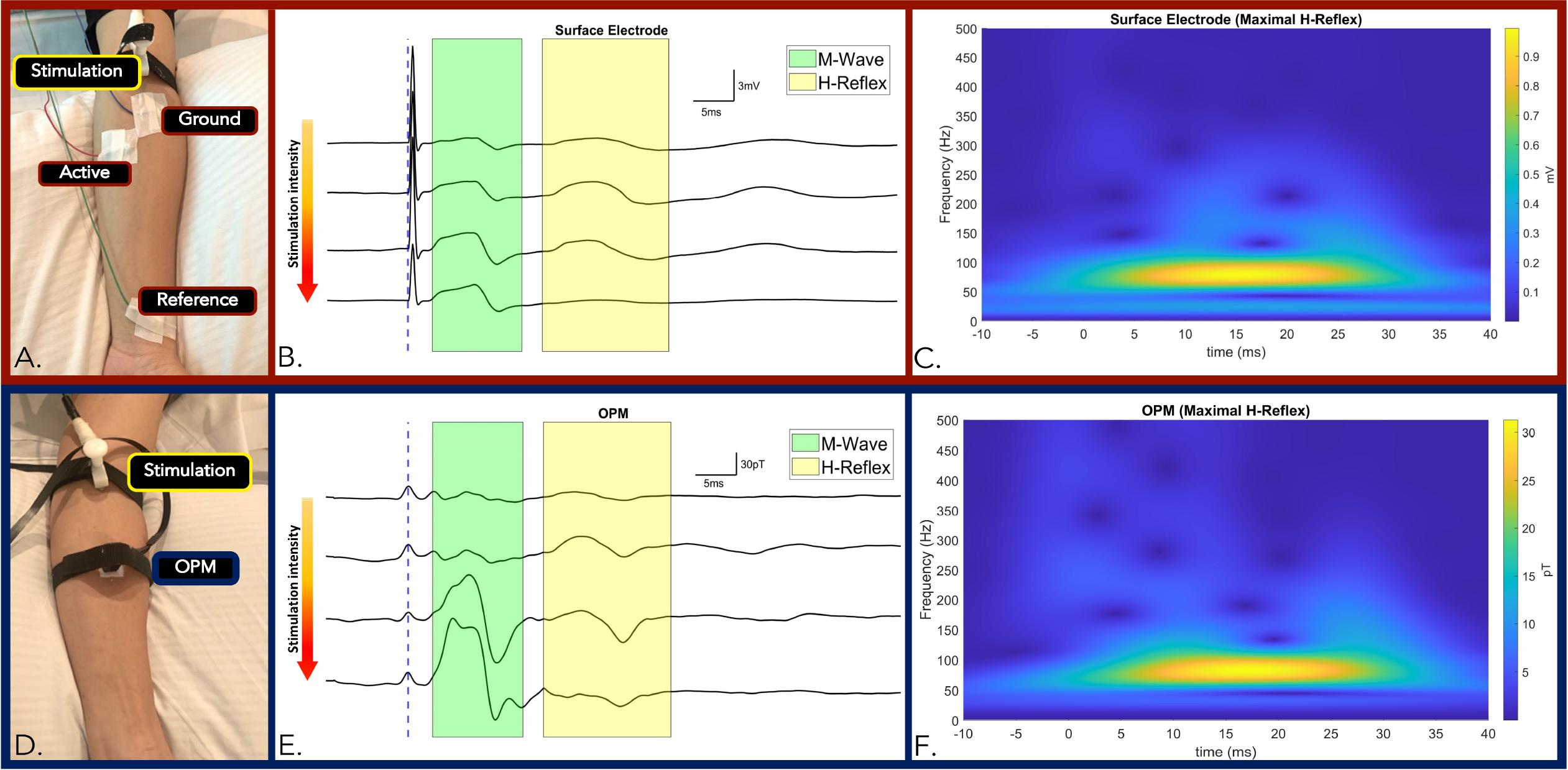
**Panel A and D:** Measurement setup of FCR M-wave and H-Reflex responses measured by surface electrode **(A)** and OPM **(D)** from the same subject. **Panel B and E:** Time-locked averages with incremental increase in stimulation intensity from top to bottom traces by surface electrode **(B)** and OPM **(E)**. The M-wave and the H-Reflex responses are highlighted green and yellow respectively. Conserved standard M and H-Reflex characteristics: 1) there was conserved incremental increases in M-wave amplitude until a maxima was reached, while 2) incremental increases in H-Reflex reach maxima, known to subsequently diminish, were observed in all measurements (**surface electrodes in B and OPM in D**). **Panel C**: Surface electrode time-frequency analysis of FCR M-wave and maximal H-Reflex recorded. **Panel D**: OPM time-frequency analysis of FCR M-wave and maximal H-Reflex recorded. (OPM = Optically Pumped Magnetometers, FCR=Flexor Carpi Radialis, mV = millivolt, pT=picoTesla, ms=milliseconds, TF=Time Frequency)

To match surface electrode time-locked measurements of the SNAP and the H-Reflex, a single OPM was positioned (with conformal Velcro strap) at the identical sites of the active electrode (at either the proximal or distal site) in the above experiments **(Figure 2E, 3D)**. In all measures, the z-direction of the OPM was adjusted normal to the skin surface in the longitudinal direction along the nerve. The y-axis of the OPM was used for measurement and the OPM were operated in single-axis mode.

To further confirm the source of the detected signals and to measure the spatial fall-off of the H-Reflex. 3 OPM sensors were placed in transverse orientation to the FCR at the one-third (proximal FCR) and one-half (distal FCR) distance between the medial epicondyle and radial styloid (**Figure 4A, 4B**). The proximal and distal OPM sensors were separated longitudinally by 4.5 cm. The central OPM sensor was placed directly over the FCR muscle. At the equipotential level, the two remaining OPMs were positioned radially 0.5 cm lateral and medial to the central OPM sensor.

**FIGURE 4.**
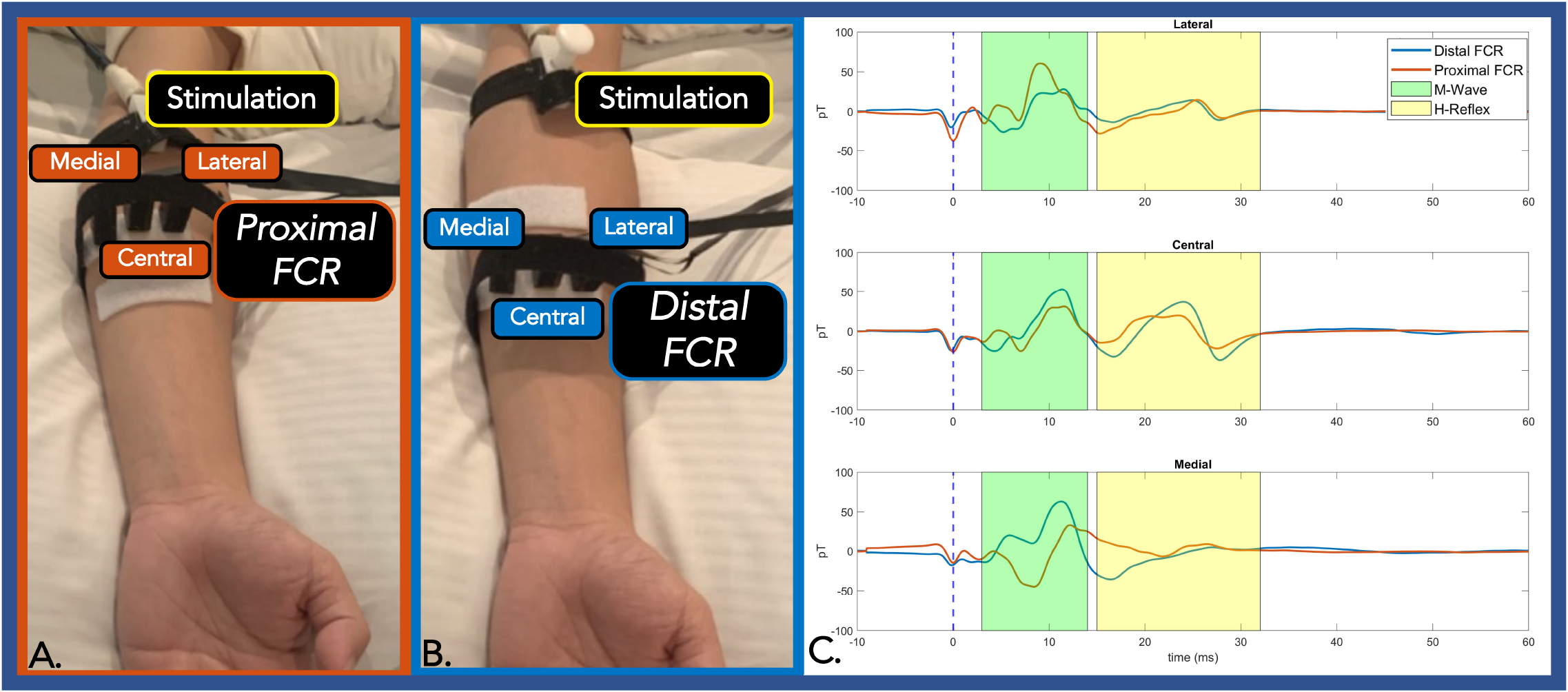
**Panel A and B**: Three OPMs were transversely placed at an equipotential level on the proximal FCR **(A)** and distal FCR **(B). Panel C:** The centrally placed OPM (directly over the FCR) exhibited the largest H-Reflex pk-pk amplitude compared with the two radially placed OPM sensors. At the proximal FCR, the lateral and the medial OPM showed reverse polarity. At the distal FCR, where the muscle spindle is thin, the medial OPM placed only detected M-Wave activity. The M-wave and the H-Reflex responses are highlighted green and yellow respectively. (OPM = Optically Pumped Magnetometers, FCR=Flexor Carpi Radialis, pT=picoTesla, ms=milliseconds).

### 2.3 Processing Pipeline

The recorded surface electrode signal was amplified and bandpass filtered from 3-1000 Hz using a Digitimer D360 isolated amplifier (Digitimer Ltd, UK) with a gain setting of 1000 for SNAP measurement and 200 for H-Reflex measurement (Christie et al., 2005). The signal was sampled by a CED Micro1401 device at 10 kHz and recorded by Signal 8.19a software (Cambridge Electronic Design, Cambridge, UK). All stimuli were repeated with a time-locked trigger to enable trial averaging while no trial rejection was necessary due to high signal to noise ratio.

Prior to OPM Neuronal recordings, each OPM was tuned and calibrated in an absolute zero magnetic field environment in the closed door MSR with the OPM gain set to 2.7 V/nT. Of note the OPM 6^th^ order hardware digital filter at 500 Hz is known to generate up to a 15 ms ringing effect post electric stimulation (**Supplemental Figure 3**). In our measurements, stimulation artifact was post-calibrated to 0 s to eliminate intrinsic OPM and analog digital converter delay. To reduce ringing effect data contamination, both stimulation artifact and subsequent ringing effect curves were regressed as a *sinc* function by non-linear least squares method and subtracted from the averaged data. A second order 20-500 Hz bandpass filtered was then applied. All post processing was carried out with MATLAB software (MathWorks Inc, MA, USA). The same processing technique was applied on all physiological OPM measurements. To compare OPM and surface electrode neural signal bandwidth, time-frequency analysis (1-500 Hz) was computed by convolving the time-locked averaged data with complex Morlets’ wavelet (Kronland-Martinet et al., 1987) as prior described by Cohen et.al. (Cohen, 2014).

## 3 Results

### 3.1 OPM Frequency response and sensitivity

The averaged OPM normalized magnitude response (over 11 chirps) decreased as frequency was incrementally increased (**Figure 1A)**. For the bandwidth between 1 Hz to 135 Hz (**Figure 1A** green shaded area), the magnitude response decreased minimally. Above 135 Hz, the magnitude response moderately decreased, and a reduced response was recorded up 500 Hz (**Figure 1A** yellow shaded area). Above 500 Hz a large reduction in response is noted as a direct effect of the Gen2 OPM hardware digital filter in the electronic module (**Figure 1A** red shaded area). Major spikes at multiples of 60 Hz in both the magnitude response and phase response plot (**Figure 1A, 1B**) are a result of the powerline noise and its harmonics.

The OPM phase response curve is approximately linear up to 500 Hz (**Figure 1B** green and yellow shaded area), indicating a relatively constant phase delay across these frequencies. A direct calculation of the average phase delay from 5 to 500 Hz is 3.77 ± 0.078 ms. In our neurophysiological measurements below, we eliminated the delay introduced by this phase response by setting the start of the stimulation artifact to time zero, so that the temporal characteristics of our OPM measurements could be directly compared to the electrode measurements.

The OPM sensitivity was divided into three segments: below 135 Hz (**Figure 1C** green shaded area), from 135 to 500 Hz (**Figure 1C** yellow shaded area), and from 500 to 700 Hz (**Figure 1C** red shaded area). Ignoring outlier frequencies caused by powerline and other technical noise sources, our measured sensitivities in the three segments are: 17.7 ± 3.5 fT/√Hz below 135 Hz, 20.8 ± 3.8 fT/√Hz from 135-500 Hz, and 101.6 ± 85.6 fT/√Hz above 500 Hz. Filtering at 60 Hz and its harmonics aimed to remove powerline noise inherently resulted in downward spikes at multiples of 60 Hz while environmental noise sources lowered sensitivity below 65 Hz (**Figure 1B**). The OPM total harmonic distortion ranged from -25 dBc to -60 dBc for measured input amplitudes between 15 and 1,500 pT.

### 3.2 Ultrasound Neural Imaging Measurements

Using high resolution B-mode ultrasound imaging, we accurately measured the median nerve circumference, area, and depth at the distal bicep and proximal bicep, The median nerve at two measurement sites was similar in circumference (1.78 cm and 1.7 cm, respectively) and area (0.22 cm^2^and 0.21 cm^2^, respectively), but was located deeper at the proximal bicep (0.64 cm) when compared to the distal bicep (0.43 cm), (**Supplemental Figure 4)**.

### 3.3 Optically Pumped Magnetometer Electrodiagnostic Measurements

#### 3.3.1 Sensory Nerve Action Potential and Current

We first verified OPM-based detection of afferent sensory nerve action current when compared to surface electrode SNAP in median nerve. We observed identical surface electrode and OPM 0.8 ms peak latency temporal dispersion **(Figure 2B, 2F)** and demonstrated nerve conduction velocity of 50 m/s largely in line with reported median nerve (large fiber) orthodromic sensory conduction velocities (Shehab, 1998;Valls-Sole et al., 2016). Additionally, surface electrode (inter-electrode distance of 6 cm) action potential (time frequency derived) center frequencies, were largely equivalent (260 Hz) when compared to OPM (200 Hz) **(Figure 2C-D, 2G-H)**. Of note, surface electrode (inter-electrode distance of 2 cm and 4 cm) action potentinal (time frequency derived) center frequencies (460 Hz and 300 Hz) were greater than the OPM center frequency **(Supplemental Figure 2)**.

The effective current density of the nerve was simulated and back-calculated by the Biot-Savart magnetic MATLAB toolbox (Loic, 2015) by employing the detected B-field action current and median-nerve-depth-to-vapor-cell distance (including the 6.5 mm sensor standoff). The nerve was simulated as a straight wire. The estimated total current was 0.195 μA and 0.19 μA from the median nerve at proximal and distal bicep, respectively. The current density was calculated as 0.92 μA/cm^2^ and 0.91 μA/cm^2^ at the proximal and distal bicep, respectively.

### 3.3.2 Flexor Carpi Radialis Hoffman-Reflex

Two experiments were conducted to verify accurate OPM-based detection of M-wave and H-Reflex of the FCR muscle. In our first experiment, stimulation intensity was gradually increased to elicit conserved standard M and H-Reflex characteristics: 1) there was conserved incremental increases in M-wave amplitude until a maxima was reached, while 2) incremental increases in H-Reflex reach maxima, known to subsequently diminish. Four demonstrative traces observed with incremental increases in stimulation intensity (Figure 3B, 3E). We compared the electromyography response and the reflex reaction of the FCR muscle between the gold standard surface electrode measures and the OPM. In subjects (N=2) we observed conserved standard M and H-Reflex characteristics with stimulation intensity increases in all measurements (OPM and surface electrodes). M-Wave (3.1 ms) and H-Reflex (17.6 ms) onset latencies were identical for surface electrode and OPM measurements **(Figure 3B, 3E, Supplemental Table 1)**. M wave and H-reflex time frequency analysis center frequency of 80 Hz was highly conserved across OPM and surface electrode measurements **(Figure 3C, 3F)** and M-wave general shape was conserved (Merletti et al., 1995).

In the second experiment, three radially placed OPMs were positioned at the one third (proximal FCR) and one half (distal FCR) distance between the medial epicondyle and radial styloid. At the proximal FCR, all three OPM successfully detected M-wave and H-Reflex activities. The central OPM had the largest H-Reflex pk-pk amplitude among the three measurements and the medial OPM showed reversed polarity. At the distal FCR, where the muscle spindle gets thinner, the central OPM measurements again showed larger H-Reflex activities compared with the lateral sensor. The medial OPM that was placed closer to the palmaris longus muscle only detected muscle activity **(Figure 4C)**.

## 4 Discussion

Widely adopted by clinical Neurophysiologists, conventional surface electrode electrodiagnostic measures (i.e., SNAP and H-Reflex) are routinely deployed to measure functional and dysfunctional neural physiology. Our work demonstrate, QuSpin Gen2 OPM can measure frequencies up to 500 Hz with a sensitivity capable of action current detection (SNAP and H-Reflex equivalent) confirmed with conventional surface electrode measures.

The presented frequency response curve and frequency dependent sensitivity (**Figure 1)** indicate that the OPM sensor can measure frequencies: 1) above the reported bandwidth (135 Hz), and 2) within range of peripheral nerve action current frequencies (up to 500 Hz), but 3) with a reduced sensitivity in this range (135-500 Hz). Generally, the sensitivity of the OPM from 0 to 500 Hz is below 30 fT/√Hz (**Figure 1C**). A 30 fT/√Hz threshold of detection is well below the magnetic nerve action current signals detected throughout the study, which were on the range of or greater than one pT (1,000 fT). As such, the evoked magnetic activities observed throughout this report are above the minimum detection amplitude of the commercially available Gen2 OPM sensor and within its current capabilities.

The OPM linear phase response (between 0 and 500 Hz) indicate preservation of the phase relationships of different frequency components below 500 Hz necessary for peripheral nerve action current detection (**Figure 1B**). The observed nominal total OPM harmonic distortion (less than -25 dBc) indicate physiologic signals do not significantly distort nerve action current measures.

After sensor characterization, OPM fast fiber sensory nerve action current detection was confirmed with conventional surface electrode measurements. Identical SNAP temporal dispersion was measured with OPM surface electrode SNAP recordings. In both sensors, temporal dispersion derived nerve conduction velocities match expected large fiber 50 m/s conduction velocity. To our knowledge, this is the first measure of peripheral nerve action current with OPM.

It should be noted that with 6 cm surface electrode separation center frequencies of the action potential and action current were largely equivalent albeit with a nominal 60 Hz difference (**Figure 2C, 2D, 2G, 2H**). But with decreased surface inter-electrode distance (4 cm and 2 cm active to reference), we observed an incrementally increased action potential width and action current center frequency difference (100 Hz at 4 cm and 260 Hz at 2 cm) when compared to OPM (**Supplemental Figure 2**). These differences are due to a combination of factors. Because it takes a finite time (0.8 ms) for a peripheral large fiber sensory nerve action potential to travel from onset to maxima, the traveling action potential spatial distribution is calculated as approximately 4 cm with a concomitant nerve conduction velocity of 50 m/s (Dumitru et al., 2010). To accurately measure the full action potential signal, a minimum of 4 cm inter-electrode distance is required. But the closer the electrodes are placed (less than 4 cm), the more of the signal will be eliminated and hence the smaller it will appear (i.e., action potential amplitude and duration) (Dumitru et al., 2010). In line, we detected a decrease in surface electrode center frequency with larger inter-electrode distances (460 Hz at 2cm, 300 Hz at 4 cm, and 260 Hz at 6 cm) **(Supplemental Figure 2, Figure 2C, 2D)**. Precisely positioned over the active electrode site, the OPM acts as a single sensor with a fixed effective distance, (i.e., circumferential diameter at which the median nerve propagating action current is detected), that likely contributes to the observed 200 Hz center frequency. In addition to the OPM fixed effective distance that potentially widened the action current, the qualitative amplitude reduction of high frequency action current components is most likely due to the OPMs intrinsic low pass filter, but other as yet identified factors may also contribute. Nonetheless, the OPM was capable of peripheral nerve action current detection (albeit with reduced response), that inevitably preserved the equivalent (0.8 ms) surface electrode derived temporal dispersion. In sum, OPM accurately measures sensory nerve action current latency, however the above-described factors may result in relative amplitude blunting due to a fixed effective distance, intrinsic sensitivity, qualitative filtering and other undetermined factors. When considering OPM sensory nerve action current measures as a clinical diagnostic adjuvant, further work is essential in clinical disease populations (i.e., axonal neuropathy) where relative amplitude comparison between upper, lower, right and left limbs is requisite.

Of note the signals from surface electrodes and OPM are similar because they fundamentally arise from the same moving source (i.e., the median nerve propagating action potential or current). However, there are important distinctions. The OPM magnetometer measures the field vector while the surface measures a scalar quantity. Inherently, more information is captured (i.e., direction of the field) with OPM when compared to electrodes, that may improve nerve action source localization with expanded N=10 OPM sensor arrays. Moreover, we observed equivalent current densities: 1) 0.92 μA/cm^2^ (0.64 cm median nerve depth at proximal position) and 2) 0.91 μA/cm^2^ (0.43 cm median nerve depth at distal position), that in aggregate evince a capability to identify depth independent median nerve action currents. In the near future, OPM depth independent current density equivalents may: 1) expedite peripheral nerve source localization and peripheral nerve differentiation critical for both neural recording as well as targeted stimulation, and 2) may supersede spatial resolution obtained with conventional electrode based finite element modeling. Clinically, (if developed) these depth independent current density algorithms may better identify neural root, trunk, division, cord, and branch dysfunction in plexopathies where multiple neural structures are in close proximity.

Analagous to SNAP measures, OPM M-wave and H-Reflex were verified when compared to gold standard surface electrode measures. Time-locked OPM derived M-wave and H-reflex responses demonstrated the same temporal latency and center frequency when compared to those from the surface electrode measurement. Additionally, as we increased the stimulation amplitude, we observed an initial increase and then decrease in the H-Reflex amplitude, highly characteristic of an H-Reflex signal(Christie et al., 2005;Stowe et al., 2008).

Next, we identified a maximal peripheral FCR H-Reflex current source by distributing three separate OPM sensors within an equipotential line at both proximal and distal FCR. At both transverse levels, the centrally placed OPM (directly over the FCR) exhibited the largest H-Reflex pk-pk amplitude compared to the two radially placed OPM sensors. At the proximal FCR site, the lateral and medial OPM showed reverse polarity, demonstrating the propagation direction and the center location of the propagating action current. Collectively, this location dependent pattern of OPM signal magnitude (i.e., largest H-Reflex action current identified with Central OPM sensor placed directly over the FCR) supports peripheral nerve action current identification with OPM multi-sensor array methodology. Reliability of conventional FCR H-Reflex identification is low (Christie et al., 2005). Based on these preliminary results, OPM arrays distributed over the forearm area (4-10 OPMs) may expedite and improve reliable and objective identification of the FCR H-Reflex response.

Similar to single sensor, multi-OPM peak latencies were measured within typical range of conventional gold standard surface electrode FCR H-Reflex latency (17-18 ms) (Jabre, 1981;Christie et al., 2005;Khosrawi et al., 2015). Future clinical cohort studies are planned in patients with known demyelinating disease, plexopathy or radiculopathy with the aim to identify pathological absent or slowed H-Reflex measured with OPM.

While our OPM action current recordings are encouraging, our work is not without limitations. First, the OPM is only operational within a MSR which limits general use in the open conventional clinic setting. Future work will focus on the development of open field capable small single limb (arm or leg) magnetic shielding devices that may afford convenient in office OPM electrodiagnostic (SNAP and H-Reflex) characterization, that inherently may lower costs when compared to conventional MSR. Second, post-hoc stimulation artifact and ringing effect removal was required to obtain early SNAP measures. Future automated, optimization of OPM filtering techniques applied to the transimpedance amplifier interaction with the lock-in amplifier are planned; this together may minimize the ringing effect. Additionally, accurate OPM neuronal action potential measurement requires informed placement of each sensor, i.e., B-mode median nerve and Flexor Carpi Radialis imaging. Employing a multiarray 4-10 OPM sensor solution has the potential to circumvent the requirement for B-mode ultrasound scanning and therefore may improve SNAP and FCR H-Reflex reliability. As prior mentioned, OPM intrinsic sensitivity and filtering may result in amplitude blunting of fast component sensory nerve action currents. Future clinical cohort studies may define OPM sensory nerve action current amplitude characteristic accuracy and reliability as a clinical electrodiagnostic.

In summary, we demonstrate two major findings: 1) the commercially available QuSpin Gen2 OPM is capable of measuring signals at a frequency above 135 Hz (up to 500 Hz) with a sensitivity and phase response appropriate for the detection of peripheral nerve action current (SNAP and H-Reflex) equivalents, and 2) OPM has comparable temporal resolution when measuring SNAP and H-Reflex equivalents consequently compared to gold standard surface electrodes. Taken together, our results warrant further OPM sensor electrodiagnostic investigation in clinical disease cohorts.

## Contribution to the field

Electrodiagnosis is routinely integrated into clinical neurophysiology practice for peripheral nerve disease diagnoses such as neuropathy, demyelinating disorders, nerve entrapment/impingement, plexopathy or radiculopathy. Well established in clinical neurophysiology practice, surface electrode electrodiagnostics routinely identify peripheral neuropathy, and nerve entrapment with the Sensory Nerve Action Potential (SNAP); whereas plexopathy, demyelinating disease and radiculopathy are identified with the Hoffmann Reflex (H-Reflex). The propagation of peripheral nerve action potentials along a peripheral nerve are the result of ionic current flow which, according to Ampere’s Law, generates a small magnetic field (action current). Termed magnetoneurography, superconducting quantum interference devices (SQUID) are capable of peripheral action current detection. Optically pumped magnetometers (OPMs) are an emerging class of quantum magnetic sensors with a demonstrated sensitivity at the 1 fT/√Hz level, capable of cortical action current detection. Based on these capabilities we hypothesized OPM may also detect peripheral nerve action current signatures. With careful OPM bandwidth characterization, we provide compelling evidence that OPMs can detect peripheral nerve action current (SNAP and H-Reflex) equivalents consequently confirmed with conventional surface electrode measures. Taken together, our results warrant further OPM sensor electrodiagnostic investigation in clinical disease cohorts.

## Ethics statements

### Studies involving animal subjects

Generated Statement: No animal studies are presented in this manuscript.

### Studies involving human subjects

Generated Statement: The studies involving human participants were reviewed and approved by UCSD IRB 171154. The patients/participants provided their written informed consent to participate in this study.

### Inclusion of identifiable human data

Generated Statement: No potentially identifiable human images or data is presented in this study.

### Data availability statement

Generated Statement: The raw data supporting the conclusions of this article will be made available by the authors, without undue reservation.

## 6 Author Contributions

IL proposed the experimental concept. IL, YB, JP, AB, and PS designed the methodology and experiments. AB, PS, RR, DK, VS, TC, MH, and IL suggested experimental improvements. IL, YB, JP performed the experiments. YB and JP processed and analysed the experimental data. AB provided key suggestions for the appropriate analysis the OPM sensor characterization data. IL, YB, and JP wrote the primary content of the paper. All other authors participated in the editing of the final manuscript.

## 8 Data Availability Statement

Data is available at the authors’ discretion upon direct request.

## Supplementary Material

### 1 Supplemental Figures

#### 1.1 Helmholtz Coils setup

**Supplemental Figure 1:**
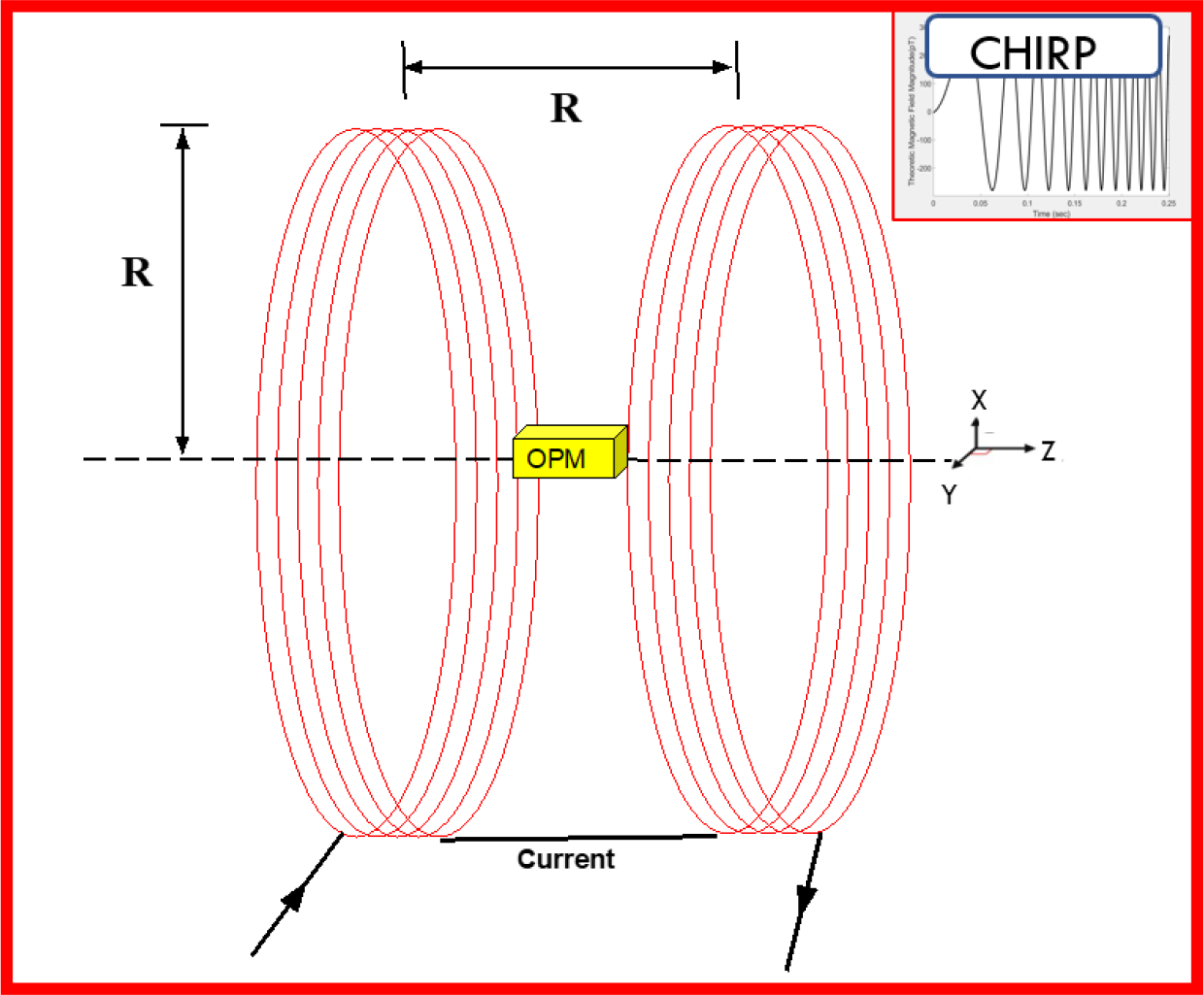
OPM frequency response and sensitivity characterization setup. A single Gen-2 OPM was placed between two copper wire Helmholtz Coils (coiled red lines are 7.5 cm radius loops with 5 coils per loop) at 7.5 cm separation. Based on electromagnetic theory, the amplitude of the magnetic field in the direction of the x-axis at the center of the two coils, is given by the equation: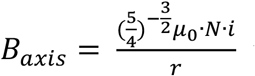, where r is the radius of the coils, μ_0_ is the relative permittivity of air, N is the number of wire loops in each coil, and i is the current running through the coils. Once the response of the OPM to several chirp functions (N=11) was recorded, the frequency response of the sensor was calculated by performing a fast Fourier transform of sensor response to each chirp and averaging each response to reduce the effects of noise. Then this Fourier transform was normalized to frequency content from the input voltage chirp. This process can be summarized mathematically with the equation: 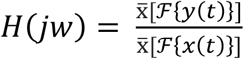,where *H*(*jw*)is the frequency response, *y*(*t*)is the signal recorded by the OPM, and *x*(*t*)is the voltage chirp function, 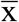 denotes calculation of the mean, and ℱ{*y*(*t*)}is the Fourier transform computed using the Fast Fourier transform algorithm. The OPM response at 5 Hz, was well within the manufactures listed bandpass of 3-125 Hz, was set to unity to create the final frequency response curve.

#### 1.2 Surface electrode SNAP measurement

**Supplemental Figure 2:**
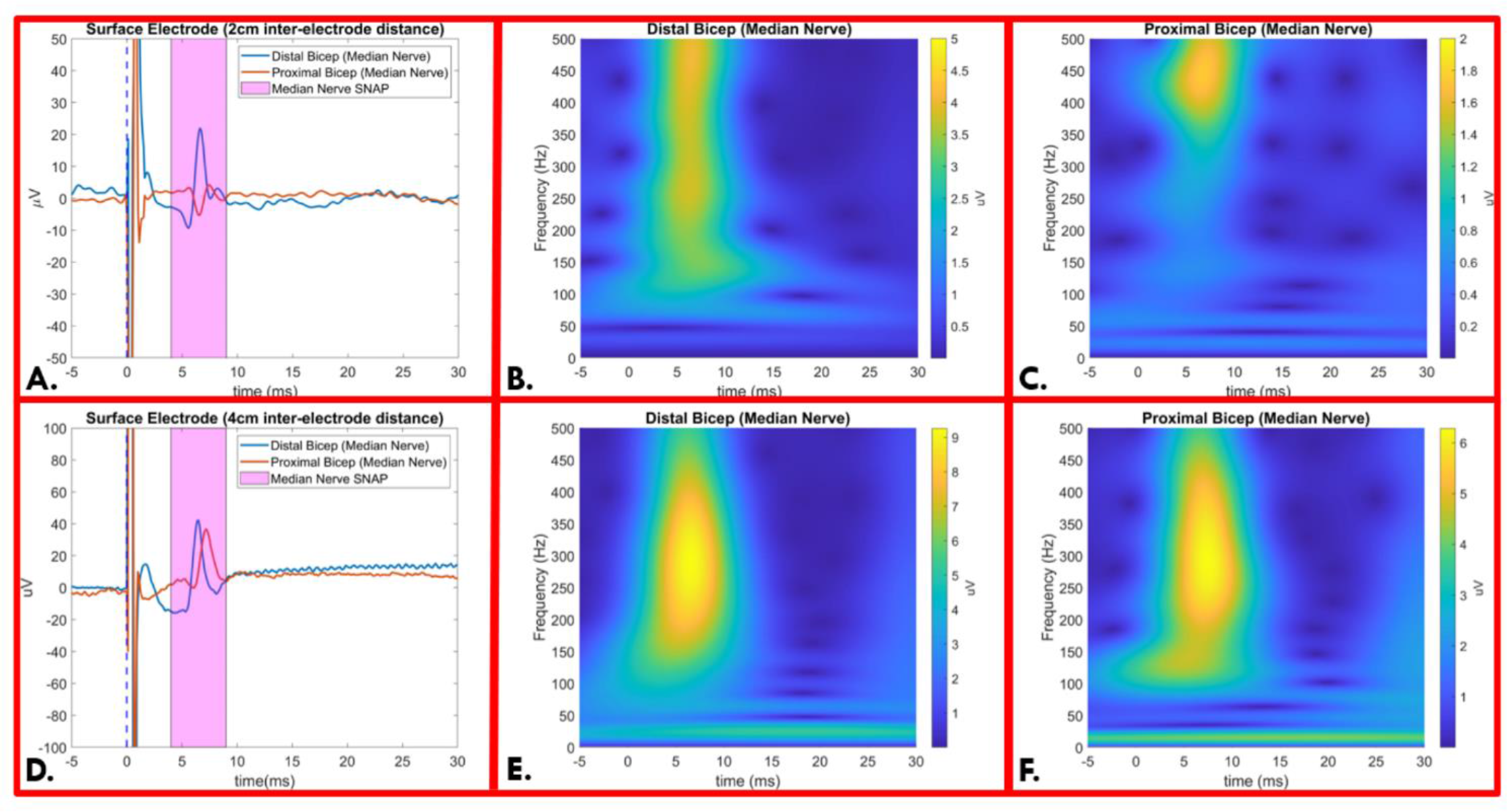
**Panel A and D:** Time-locked average comparison from surface electrode with 2 cm **(A)** and 4 cm **(D)** inter-electrode distance for median nerve SNAP measurement demonstrated 0.8 ms temporal dispersion. SNAP action potential is marked in the magenta shaded area. **Panel B and C**: 2cm inter-electrode distance time-frequency analysis for SNAP measured at distal bicep **(B)** and proximal bicep **(C)**. Distal bicep (**B**) and proximal bicep (**C**) demonstrate equal center frequencies of 460 Hz. Distal bicep residual low frequency components (**C**) are due to triphasic action potential measured from the superficial median nerve. **Panel E and F**: 4cm inter-electrode distance time-frequency analysis for SNAP measured at distal bicep **(E)** and proximal bicep **(F)**. Distal bicep (**E**) and proximal bicep (**F**) demonstrate equal center frequencies of 300 Hz. (SNAP= Sensory Nerve Action Potential, μV = microvolt, ms=milliseconds)

#### 1.3 OPM SNAP Raw Measurement

**Supplemental Figure 3:**
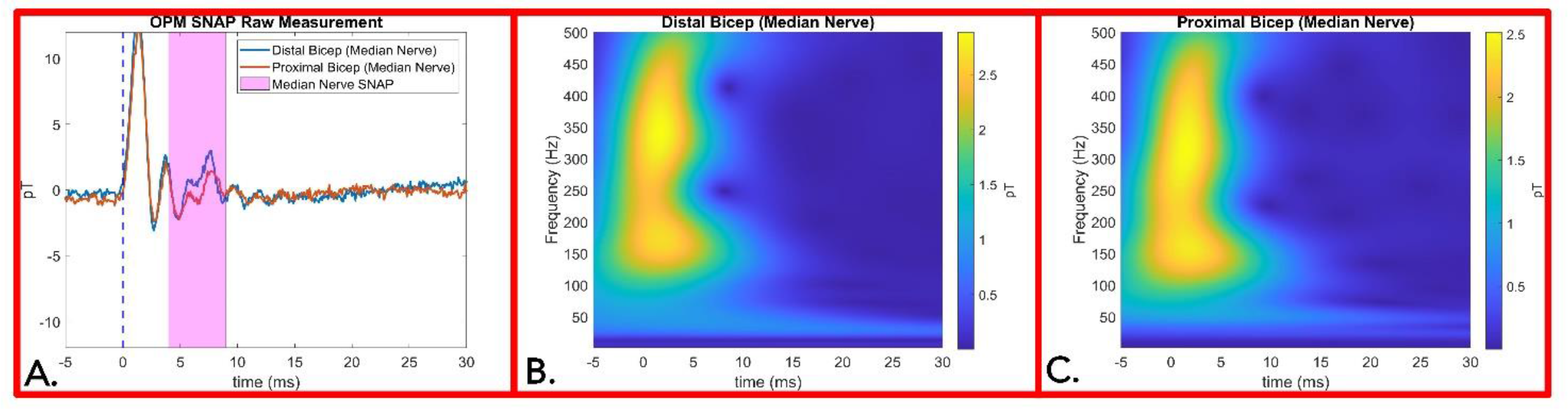
**Panel A**: OPM SNAP raw measurements from median nerve at distal bicep and proximal bicep. The OPM internal hardware digital filter at 500 Hz generate up to a 15 ms ring effect post stimulation. **Panel B and C**: Time-frequency analysis for SNAP measured by OPM before artifact removal at distal bicep **(B)** and proximal bicep **(C)**. The stimulation artifact and its ringing effect distort accurate signal acquisition of the SNAP center frequency. To reduce ringing effect data contamination, both stimulation artifact and subsequent ringing effect curves were regressed as a *sinc* function by non-linear least squares method and subtracted from the averaged data. Finally, a second order 20-500 Hz bandpass filtered was applied effectively removing stimulation artifact ring effects **presented in Figure 2F**. (OPM = Optically Pumped Magnetometers, SNAP= Sensory Nerve Action Potential, pT=picoTesla, ms=milliseconds)

#### 1.4 Ultrasound Neural Imaging Measurements

**Supplemental Figure 4:**
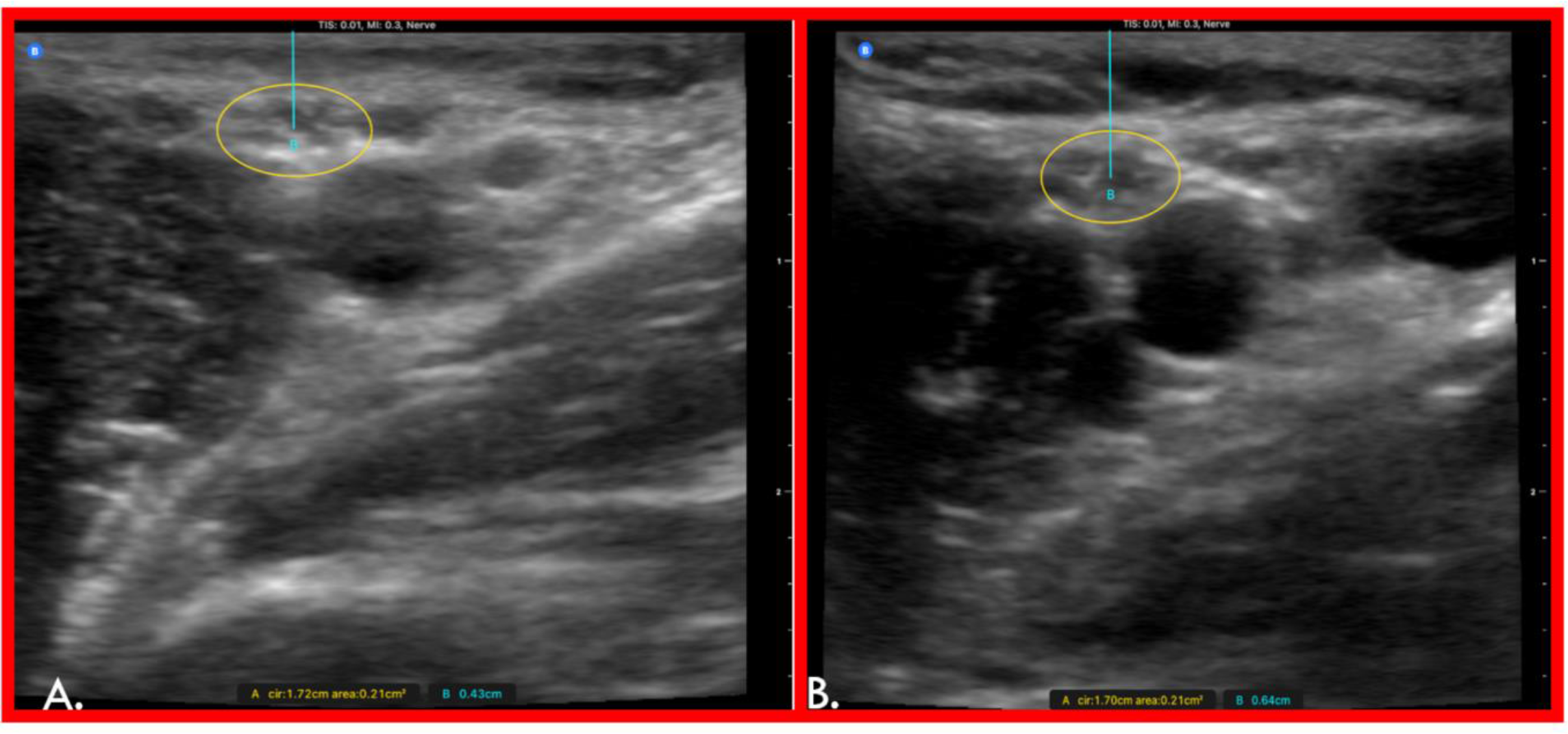
Using high resolution B-mode ultrasound imaging, we accurately measured the median nerve circumference, area, and depth at the distal (**Panel A)** and proximal (**Panel B)** bicep. The median nerve at two measurement sites was similar in circumference (1.72 cm and 1.7 cm, respectively) and area (0.21 cm^2^ for both) (**Panel A & B**). Compared to the median nerve localized at the distal bicep (0.43 cm) (**Panel A**) it was localized deeper at the proximal bicep (0.64 cm) (**Panel B**).

### 2 Supplemental Table

**Supplemental Table 1:**
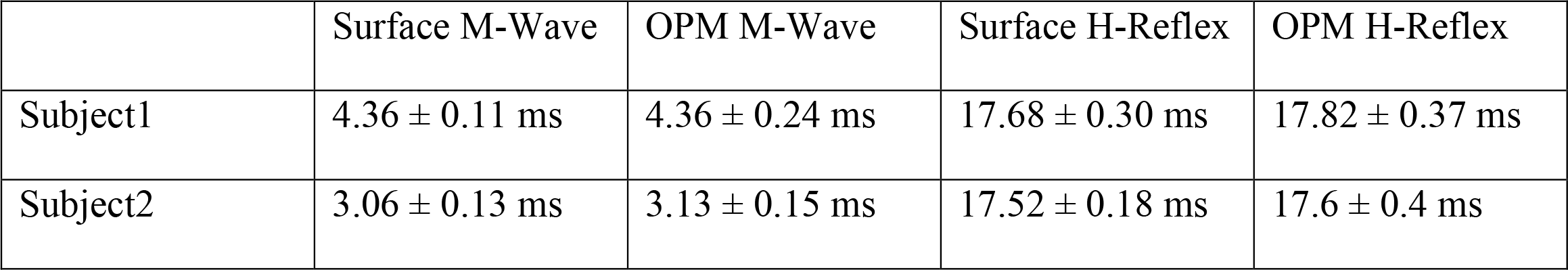
Onset latency statistic comparison for M-wave and H-Reflex measurements between Surface electrode and OPM from two subjects.

|  | Surface M-Wave | OPM M-Wave | Surface H-Reflex | OPM H-Reflex |
| --- | --- | --- | --- | --- |
| Subject1 | 4.36 ± 0.11 ms | 4.36 ± 0.24 ms | 17.68 ± 0.30 ms | 17.82 ± 0.37 ms |
| Subject2 | 3.06 ± 0.13 ms | 3.13 ± 0.15 ms | 17.52 ± 0.18 ms | 17.6 ± 0.4 ms |

